# Simulation of cell-size systems at long timescales with flexible protein structures

**DOI:** 10.64898/2026.06.20.733545

**Authors:** Kamila Yunas, Amar Singh, Matthew M. Copeland, Andrii M. Tytarenko, Petras J. Kundrotas, Randal Halfmann, Pavlo O. Kasyanov, Eugene A. Feinberg, Ilya A. Vakser

## Abstract

Protein behavior inside cells is dominated by the crowded nature of the intracellular environment. Progress in structure determination of proteins and protein complexes, based on advances in Artificial Intelligence, provides an opportunity for structure-based modeling of cellular phenomena. Such modeling at the atomic resolution has been advanced by the traditional simulation techniques, e.g. molecular dynamics. A recently developed docking-based approach implements Markov Chain Monte Carlo sampling of intermolecular energy landscapes, offering several orders of magnitude faster simulation protocols. The approach allows addressing much longer trajectories of macromolecular systems in the crowded intracellular environment at atomic resolution. The sampling by design avoids low-probability (high-energy) states, which greatly accelerates the simulation process. A notable feature of this docking-based approach is the rigid body approximation of protein structures. The rigid-body approximation had been the primary direction in the protein docking field up until recent developments in deep learning. The rigid-body approach should be quite robust for the higher energy transient interactions that dominate the highly crowded cellular environment, as they likely involve relatively small conformational change. However, it is less applicable to the low-energy protein-protein complexes, especially those involving flexible regions. We addressed this problem by incorporating AlphaFold3 top models of the protein complexes in the mapping of the intermolecular energy landscape, as representative of the low-energy configurations of the protein assembly. By the nature of the AlphaFold predictions, these models involve appropriate conformational change between unbound and bound structures. These low-energy docking poses are combined with the rigid-body docking predictions that cover the multiplicity of the transient interactions. Such combination directly addresses the conformational flexibility of proteins upon binding along with the multiplicity of the transient protein encounters in the crowded cellular environment.

**SIGNIFICANCE:** Protein behavior inside cells is dominated by the crowded nature of intracellular environment. A recently developed approach allowed addressing long simulation trajectories of macromolecular systems in such environment at atomic resolution. A notable feature of this approach is the rigid body approximation in representation of the protein structures, which had been popular in the field up until the recent developments in artificial intelligence. However, such approximation is less applicable to stable protein-protein complexes, especially those involving flexible regions. We addressed this problem head-on by incorporating top deep learning-generated models of protein complexes. The new approach directly accounts for the flexibility of protein structures upon binding, along with the multiplicity of the transient protein encounters in the crowded cellular environment.

## INTRODUCTION

Molecular mechanisms of cellular processes are largely based on protein interactions. Our ability to understand and modulate physiological processes benefits from the knowledge of structural details of proteins and the kinetics of their interaction (1,2). Protein behavior inside cells is dominated by the crowded nature of the intracellular environment (3). Spectacular progress in structure determination of proteins and protein complexes, based on the advances in Artificial Intelligence (4-6), provides an opportunity for structure-based modeling of the cellular phenomena. Such modeling at the atomic resolution has been advanced by the traditional simulation techniques, e.g. molecular dynamics. A recently developed alternative approach based on Markov Chain Monte Carlo (MCMC) sampling of the intermolecular energy landscapes offers several orders of magnitude faster procedures, allowing us to address much longer trajectories of macromolecular systems in the crowded intracellular environment at atomic resolution (7,8). A systematic protein docking Fast Fourier Transform (FFT)-based procedure (9-11) places minima on the intermolecular energy landscape, subsequently sampled by the MCMC protocol, in which a protein moves from one assembly to another, with moves accepted or rejected according to the Metropolis criterion. The sampling by design avoids low-probability (high-energy) states, which greatly accelerates the simulation process, thus allowing addressing physiological processes far beyond the scope of the alternative atomic resolution techniques.

A notable feature of this approach is the rigid body approximation in representation of the protein structures, which had been the primary direction in the protein docking field (9,12,13) up until the recent developments in deep learning (4,14). The rigid-body docking is robust because of a generally limited scope of conformational change upon protein binding (15), except cases of disordered regions/proteins. Also, the higher energy transient interactions, which dominate the highly crowded cellular environment should involve relatively small conformational change. Still, conformational change upon protein binding plays an essential role in protein interactions (16,17), and the rigid-body approximation, in general, is less applicable to the low-energy protein-protein complexes, especially those involving flexible regions.

We addressed this problem head-on by directly including the AlphaFold3 (AF) top models of the protein-protein complexes in the mapping of the intermolecular energy landscape, as representative of the low-energy configurations of the protein-protein assembly. By the nature of AF predictions, these models involve appropriate conformational change between unbound and bound structures. The low-energy docking poses are combined with the rigid-body FFT docking predictions that cover the multiplicity of the transient interactions. Such combination directly addresses the conformational flexibility of proteins upon binding, along with the multiplicity of the transient protein encounters in the crowded cellular environment.

## METHODS

### Basic Simulation Protocol

In the simulation protocol the state of the system consists of the molecules’ coordinates and the energy. A new state is sampled given the current state, following the MCMC paradigm. The intermolecular energy landscape of the system is represented by 30,000 lowest-energy GRAMM docking predictions for each binary protein-protein combination (9,11), corresponding to van der Waals (vdW) potential (10) or the AACE energy (18) (30,000 lowest AACE energy predictions rescored from the top 300,000 vdW predictions).

Initially, proteins are placed randomly on a grid. The number of protein copies is calculated according to the preset protein volume fraction. The position of each protein is described by the 3 × 3 rotation matrix and the translation vector relative to the origin of the coordinates. The Monte Carlo (MC) move is initiated by a random selection of a protein (“ligand”) considered for a move to other proteins (“receptors”). The receptor to move to is selected randomly among all proteins within 50 Å radius from the ligand’s geometric center. Once the ligand and the receptor are selected, the move is chosen randomly from the precalculated lowest energy docking matches for that ligand-receptor pair.

The move is accepted or rejected according to the Metropolis criterion with the detailed balance condition (probability of accessing state *j* from state *i* has to be the same as probability of accessing state *i* from *j*). The Metropolis criterion is normalized accordingly (19) as

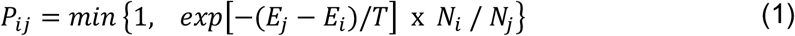

where *P*_*ij*_ is the probability of the move from step *i* to step *j*; *N*_*m*_ is the number of possible moves (receptors to move to) from state *m* with probability to be selected 1/*N*_*m*_ (*N*_*m*_ includes rejected ligand moves colliding with proteins in complex with the receptor); *E*_*m*_ is the energy of the state *m*; and *T* is the temperature of the system (a scaling factor, calibrated as *T* = 100 for vdW potential, and *T* = 4 for the AACE forcefield, following our methodology (7)). The simulation step was calibrated as 20 *ns* for vdW and 4 *ns* for AACE by matching the mean-square deviation of molecules to that simulated by molecular dynamics on the same molecular system (7). The simulation protocol implements periodic boundary condition. Among simulation observables are mean square deviation (MSD), the slope of which over time determines the rate of the protein’s translational diffusion, and residence time - the time a protein stays in a complex, calculated based on the MC move acceptance rate averaged over all copies of the protein (7).

The protocol was systematically validated on the available observations from experiments and molecular dynamics simulations. It performed consistently across different systems of proteins, at a broad range of volume fractions, in excellent agreement with experimental and theoretical results, as described in our earlier publications (7,8,20). The procedure is implemented in a publicly available GRAMMCell webserver (21).

### Conformational Flexibility

Conformational flexibility in the simulation protocol was effectively achieved by incorporating the AF predictions into the mapping of the energy landscape. The AF predictions of the structure of protein complexes start from the sequence of the participating proteins. Thus, they by design implement the flexible docking paradigm, generating bound protein conformations.

For each protein-protein complex, to map the low-energy matches involving larger conformational changes, a set of structures was generated by the AF protocol (4). These structures were added to the high-energy, transient protein matches generated by the rigid-body FFT docking implemented in GRAMM (11) (Figure 1). The AF approach by design aims at the low-energy predictions only and is not trained to predict the multiplicity of the transient protein encounters. Moreover, importantly, the transient/high-energy matches are likely to involve limited conformational changes, that are well within the accuracy of the rigid-body docking approximation.

**Figure 1.**
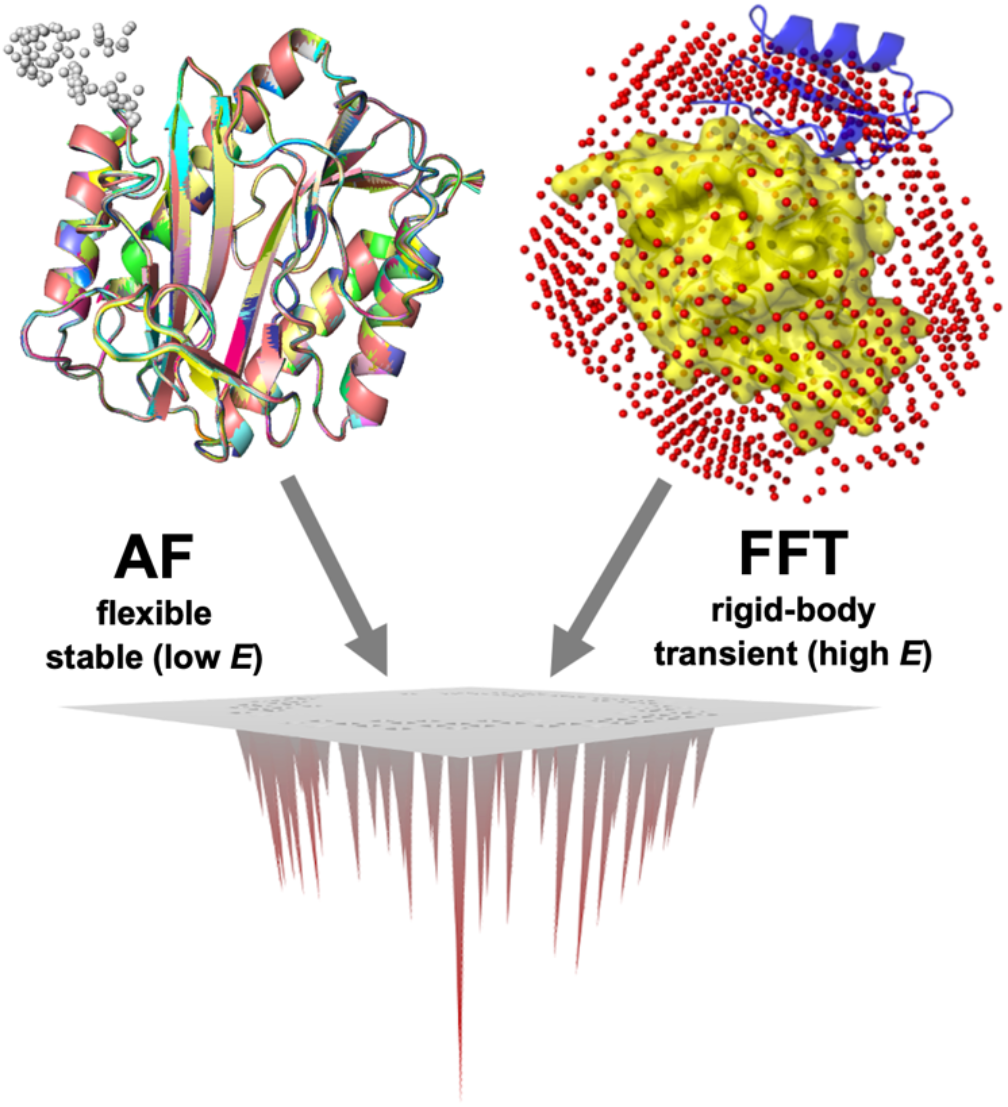
The conformational flexibility paradigm. The combination of low-energy, stable protein-protein matches by AF with the multiple transient/high-energy matches by FFT accounts for the full spectrum of protein-protein associations in the crowded cellular environment.

All predicted matches were scored by the same force field (either vdW or AACE). To account for the low-energy nature of the AF predictions, a Δ*E* value was subtracted from their energy, subject to calibration (see Results).

The coordinates of the AF predicted matches were transformed into the landscape mapping format (translation and rotation from the original coordinates to the predicted ones in the complex) readable by the GRAMMCell simulation procedure. The sampling of the landscape is performed by moving proteins in the original conformation to the AF and FFT predicted positions with the energy values from the AF and FFT modeled configurations of the complex correspondingly.

## RESULTS

### Calibration

The AF predictions by design are targeted to prediction of a few (ideally single) most stable, lowest energy structures of proteins and protein complexes, not a spectrum of docking possibilities. Thus, to combine the AF predictions with the FFT generated poses of transient encounters, we used empirical numbers for both - 100 matches modeled by AF and 30,000 matches predicted by the rigid-body FFT.

To represent the binding funnel, Δ*E* values were subtracted from the energies corresponding to the AF predictions. Different values for Δ*E* were evaluated on the simulation characteristics in the 5-mix protein set (7) at physiological concentration corresponding to the volume fraction 0.3 (Figure 2). The optimal choices were Δ*E* = 200 for vdW and Δ*E* = 8 for AACE, as the maximal values that still kept the energies of AF models comparable to the FFT energies and did not introduce bias in the energy of the system. These values did not change the observables (diffusion coefficient, aggregation number and residence time) across the range of volume fractions *V* = 0.1, 0.2 and 0.3 for both vdW and AACE potentials.

**Figure 2.**
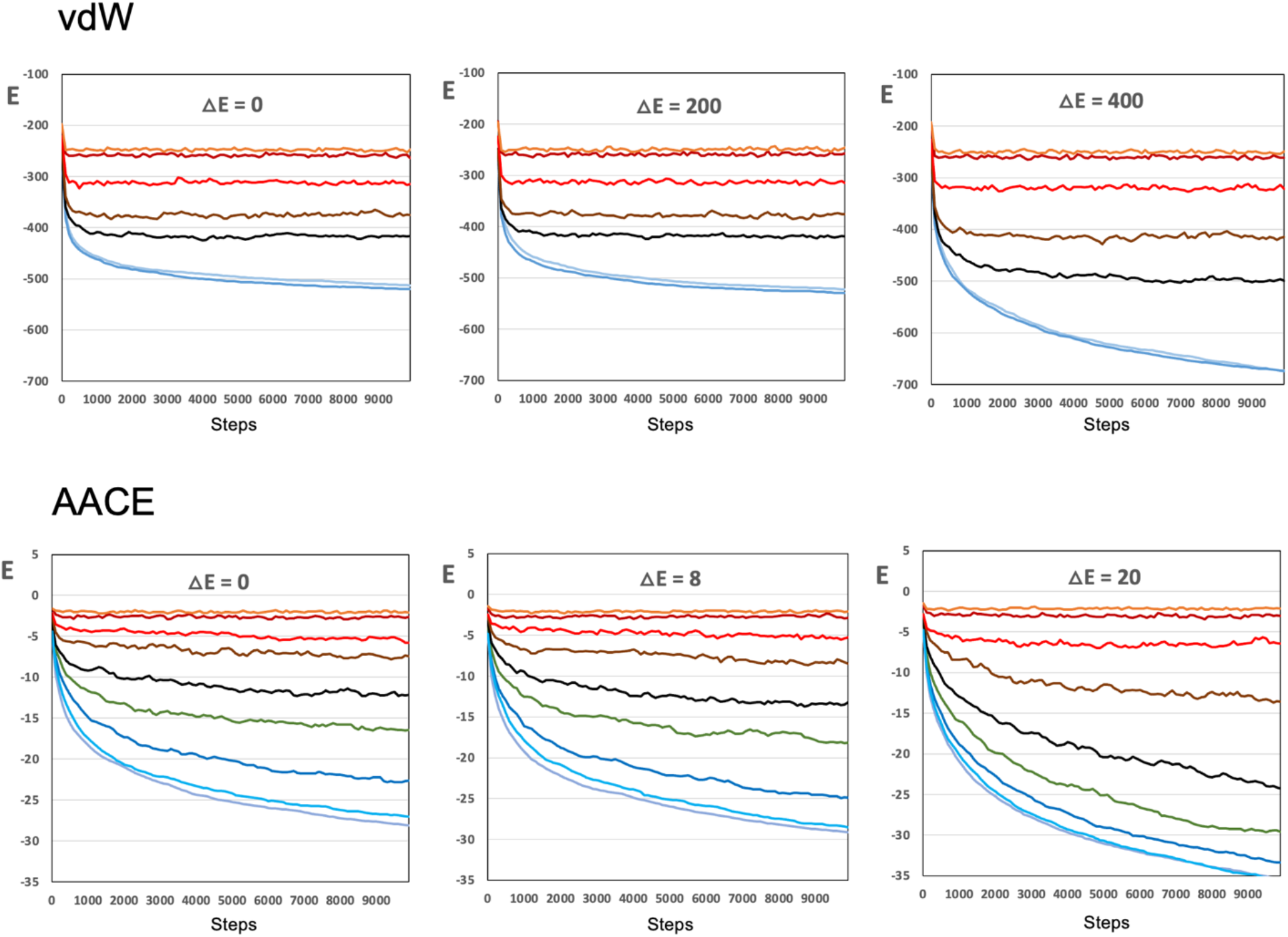
Calibration of the intermolecular energy landscape combining AF and FFT docking models. Simulations of the 5-mix set at the physiological volume fraction 0.3 and a range of temperatures ran in a 500 Å cubic box for 10,000 steps. The plots show the energy *E* of the system versus simulation steps. The data on the plots was smoothed by a 100-step averaging sliding window. The temperatures *T* = 1 to 10,000 are shown by different colors. At low temperatures (blue), the system is frozen (little or no movement of the proteins). At high temperatures (red), the system is overheated (moves accepted regardless of the energy). For each protein-protein pair, all docking poses were scored by vdW potential (top panels) or AACE potential (bottom panels). The △*E* values were subtracted from the energies of the AF predicted poses. The maximal △*E* values that still kept the energies of the AF models comparable to the FFT energies, thus not introducing bias in the energy of the system, were △*E* = 200 for vdW and △*E* = 8 for AACE potentials.

An example of the AF prediction in Figure 3 shows a conformational change at the interface between the bound (AF-predicted) and unbound proteins from *Mycoplasma genitalium* (22).

**Figure 3.**
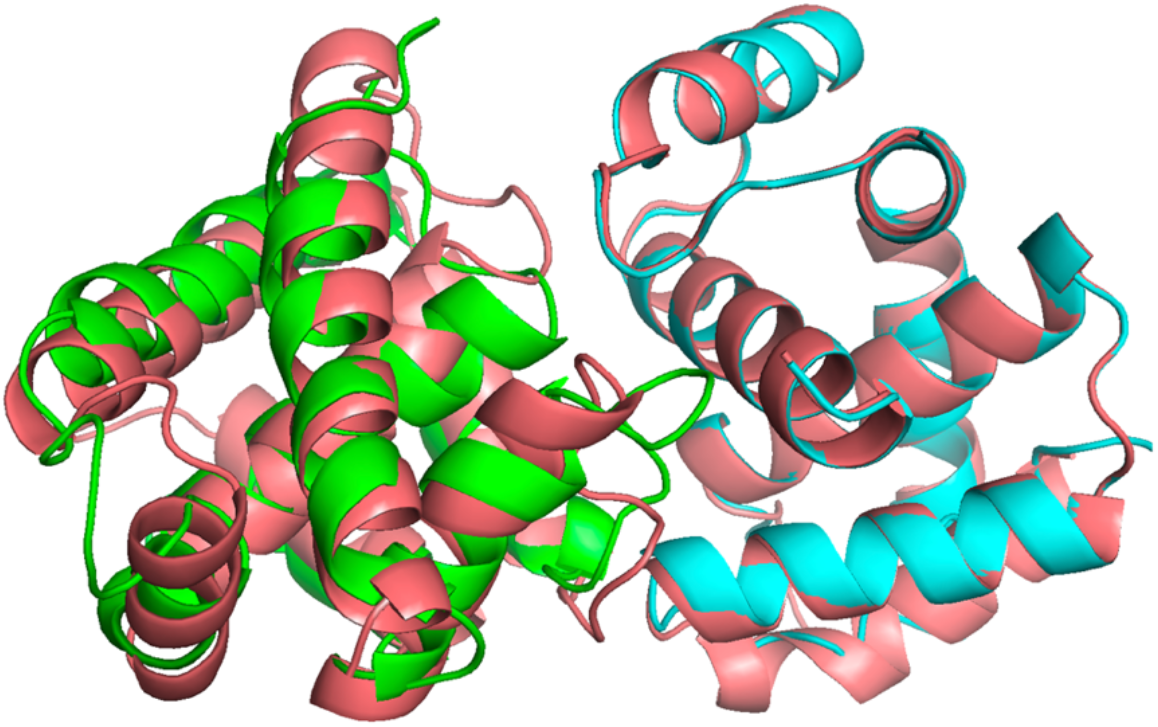
Example of conformational change incorporated in simulation. The bound structures of two monomeric proteins from *Mycoplasma genitalium* modeled by AF (magenta) are overlapped with the experimentally determined unbound structures of the corresponding proteins (1tm9 in green and 1q8c in cyan).

### Simulation of lens α-crystallins

Crystallins are the major structural proteins in the eye lens. The three main classes of crystallins - alpha, beta, and gamma - collectively account for ∼ 90% of the total protein content in the lens. The most abundant of these are the α-crystallins (Figure 4A), which belong to the small heat-shock protein (sHSP) family and function as chaperones that inhibit the aggregation of denatured or misfolded proteins, helping to prevent cataract formation. They consist of two related subtypes, αA-crystallin and αB-crystallin.

**Figure 4.**
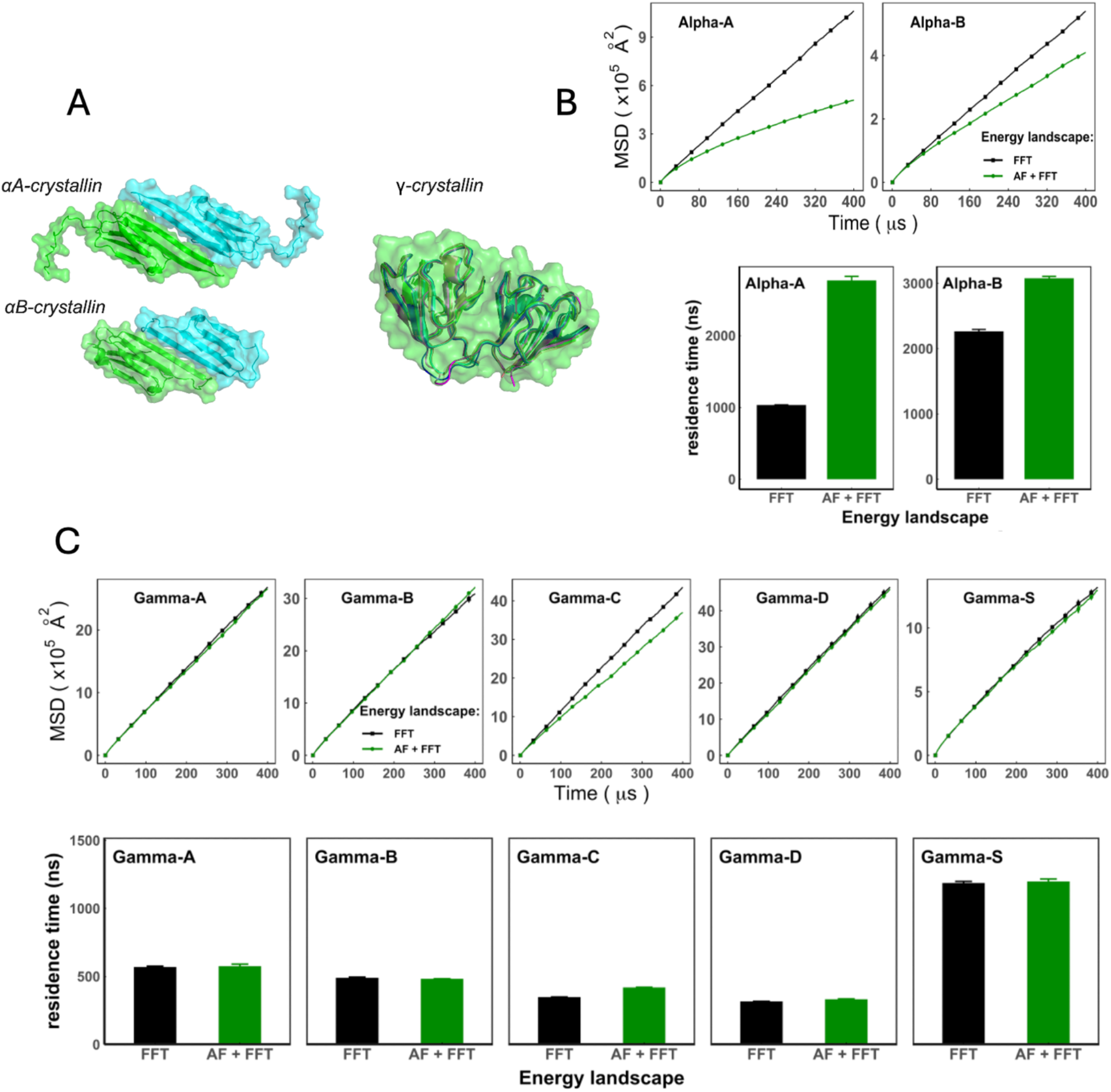
Comparison of crystallins behavior in flexible and rigid-body approaches. (A) The α-crystallins comprise two distinct proteins, αA-crystallin (3l1e) and αB-crystallin (2wj7), each assembling into oligomers containing multiple identical or highly similar chains. The γ-crystallin family (AlphaFold 3 model shown) has multiple isoforms that are structurally similar and share the same fold. (B) The AF predictions reflecting conformational changes upon binding, emphasize the known aggregation behavior of α-crystallins in comparison with the rigid-body approach (FFT). (C) The γ-crystallin is known to not form stable homo-oligomers. The lack of low-energy stable complexes is reflected in the comparable simulation results of the flexible (AF + FFT) and rigid-body (FFT) approaches.

The α-crystallins form large oligomeric assemblies. Their ordered C-terminal domain forms a stable dimeric interface, while their flexible *N*-terminal regions interact to form higher-order assemblies. There are two distinct PDB structures of the αB-crystallin oligomer (2ygd and 3j07) and one for the αA-crystallin oligomer (6t1r). Since these assemblies do not adopt a single well-defined quaternary structure, computational methods such as AF have limited ability to accurately predict their full oligomeric conformation. However, the conserved α-crystallin domain responsible for dimer formation is stable, well-structured and can be modeled using AF.

The γ-crystallins (Figure 4A) are relatively small proteins, composed of two homologous domains (*N* and *C* terminal) composed of similar motifs and connected by a short peptide. Each motif is ∼40 amino acids long, folded in a distinctive Greek key pattern. The γ-crystallin family has multiple isoforms, each coded by distinct genes, *e*.*g*., γA-crystallin, γB-crystallin, γC-crystallin, γD-crystallin, and γS-crystallin. The stability of the γ-crystallin is crucial for cataract formation, with decreased stability contributing to the development of cataracts. Unlike other crystallins, γ- crystallins do not oligomerize under normal physiological conditions.

The sequences of crystallins were obtained from UniProt entries P02489 (αA), P02511 (αB), P11844 (γA), P07316 (γB), P07315 (γC), P07320 (γD), and P22914 (γS). A total of 100 homo-dimeric protein-protein complex models were predicted using AlphaFold3. Only models with stable structures, defined by per-residue pLDDT score >70, were retained and assigned AACE energy values. Transient interactions were generated by FFT docking, as described in Methods.

The simulations were performed at physiological concentration (excluded volume fraction *V* = 0.3). Copies of each crystallin type were randomly placed in a 500 Å cubic box. Simulations were conducted for 100,000 steps (400 µs) - see Methods.

The results (Figures 4B and 4C) show the difference between the rigid body (FFT-only) and the flexible docking (AF + FFT) approaches. Stable, low-energy interactions of α-crystallins corresponding to the AF predictions, increase the residence time of the oligomers and reduce their translational diffusion (Figure 4B), thus providing a more accurate characterization of the α-crystallins behavior than the rigid-body approach alone.

The γ-crystallins are monomeric proteins and, unlike α-crystallins, do not form low-energy oligomeric complexes under physiological conditions. This is reflected in shorter residence times than those of the α-crystallins. Since there are no energetically stable interactions, as expected, inclusion of the AF-energy landscape does not change the diffusion or the residence time for the γ-crystallins (Figure 4C).

## DISCUSSION

The rigid body approximation has been a mainstream direction in the protein docking field, since its early days (9,12,13) up until the recent advance of the AlphaFold (4,14). The reason for the robustness of this approximation is a relatively small scope of conformational change upon protein-protein association in most cases (15), not involving intrinsically disordered regions/proteins. Moreover, it is reasonable to assume that transient interactions - brief protein-protein high-energy encounters that dominate the highly crowded molecular environment inside cells - involve very limited conformational change that, generally, should be well within the scope of the medium resolution FFT docking (11) used for mapping of the intermolecular energy landscape.

Still, the rigid-body approximation, in general, is less applicable to the low-energy protein-protein complexes, especially those involving flexible regions. There are different directions that potentially can be explored for inclusion of conformational flexibility into the docking-mapped MCMC-sampled simulation paradigm. For example, the problem can be directly addressed by including multiple pre-calculated conformers of the participating proteins in the docking/sampling protocol. Such approach, however, would significantly extend the compute time due to the large increase of the number of sampling points on the energy landscape, thus shortening the feasible simulation trajectories, and consequently the scope of the cellular mechanisms that could be addressed. It would also not account for the higher accuracy of the recent AF predictions of the structure of protein-protein complexes.

The essence of our approach is incorporation of the bound structures of interacting proteins. While in this study such structures were modeled by AF, they can be generated by other approaches, computational or experimental.

One limitation of the approach is that the sampling of the intermolecular energy landscape is performed by moving proteins in the original (unbound) conformation for both AF and FFT predicted positions. The energy and the docking pose of these moves to the AF-predicted matches are represented exactly by the values pre-calculated for the bound structures. However, the check for collisions with the crowders and other proteins bound to the receptor is approximate, as performed on the tentative move of the unbound structure. While this approximation is valid for most proteins, it may not be applicable to intrinsically disordered proteins with a drastic unbound/bound change in extended conformations.

## CONCLUSIONS

Protein interactions are the basis of most biomolecular mechanisms. These interactions are greatly affected by the crowded nature of the environment inside cells. Recent progress in macromolecular structure determination provided the foundation for the rapid advances in structure-based modeling of the cellular processes. The recently developed approach to simulation of protein behavior in the crowded cell-size systems implements docking-based Markov Chain Monte Carlo sampling of the intermolecular energy landscapes. The sampling is designed to avoid the low-probability energy states, thus greatly accelerating the simulation procedure, allowing addressing extra-long simulation trajectories at atomic resolution, with high accuracy inherent to the docking protocols. The approach is built on the rigid-body docking paradigm, which dominated the protein docking field up until the recent advancement of the artificial intelligence techniques. The rigid-body approach in general has been a valid approximation for structural characterization of protein-protein complexes because of the limited scope of protein bound/unbound conformational changes, especially for the high-energy transient interactions dominating the crowded environment inside cells. The rigid-body docking is less applicable to the low-energy protein interactions, especially those involving flexible fragments. We addressed the problem of conformational flexibility in the simulation protocols by incorporating the AlphaFold3 models of the protein complexes in the energy landscape mapping. Such models by design involve conformational changes upon binding. The low-energy poses predicted by AlphaFold were combined with the rigid-body docking predictions, thus covering the full spectrum of the docking poses - from the few low-energy stable assemblies to multiple high-energy transient protein encounters. The simulation procedure based on the flexible docking approach was applied to the crowded system of crystallin proteins, providing results with a closer match to the experimental observations than the rigid-body protocol.

## DATA AND CODE AVAILABILITY

The simulation procedure based on FFT docking is available through GRAMMCell resource user-friendly interface at https://grammcell.compbio.ku.edu. The AlphaFold models were generated by the AlphaFold3 pipeline with the default parameters and the number of resulting models set to 100. The code for transforming coordinates of the AF predicted matches into the landscape mapping format (translation/rotation from the original coordinates) is publicly accessible on GitHub (https://github.com/VakserLab/c2res).

## ACKNOWLEDGMENTS

This study was supported by NIH grants R35GM156453 and NSF grant DBI2224122, including IMPRESS-U supplemental funding (A.S., M.M.C., P.J.K., I.A.V.), and IMPRESS-U project 7131 (A.M.T., P.O.K). The authors wish to thank Nick Grishin for helpful discussions.

## AUTHOR CONTRIBUTIONS

I.A.V. and P.J.K. designed research, R.H. provided data, K.Y., A.S., P.J.K. and A.M.T. performed research, M.M.C. assisted in software development and running, P.J.K., E.A.F. and P.O.K. analyzed the results, I.A.V. wrote the manuscript.

